# Sequence Entropy and the Absolute Rate of Amino Acid Substitutions

**DOI:** 10.1101/056325

**Authors:** Richard A. Goldstein, David D. Pollock

## Abstract

The evolution of model proteins under selection for thermodynamic stability suggests parallels between evolutionary behavior and chemical reaction kinetics. We developed a statistical mechanics theory of protein evolution by dividing amino acid interactions into site-specific and ‘bath’ components, and show that substitutions between two amino acids occur when their site-specific contributions to stability are nearly identical. Fluctuating epistatic interactions drive stabilities into and out of these regions of near neutrality, with the time spent in the neutral region and thus the rate of substitution governed by physicochemical similarities between the amino acids. We derive a theoretical framework for how site-specific stabilities are determined, and demonstrate that substitution rates and the magnitude of the evolutionary Stokes shift can be predicted from biophysics and the effect of sequence entropy alone. Population genetics underlays our analysis, but population size does not determine the absolute rate of amino acid substitutions.

## Introduction

Modeling the rate at which protein sequences change is central to understanding how proteins adapt to their structural, functional, and thermodynamic requirements. It is also key to deciphering the patterns of conservation and variation that reflect evolutionary processes and the properties of specific proteins. An important step was Kimura’s calculation of the probability of fixation of a single mutation given constant relative fitnesses of the wild type and mutant (*1–3*). Fixation probabilities alone, however, do not address how or why fitness differences come to be, and therefore cannot explain observed substitution rates. Empirically derived substitution rates have long been obtained by analyzing differences between related protein sequences (*4–6*), providing estimates of average rates but not explaining them. Although this approach has been extremely useful, its successes were achieved by ignoring the underlying biophysics, molecular biology, and population dynamics, as well as how these rates vary amongst sites and time.

In recent years, the number of protein sequences, computational speeds, and our knowledge of protein biophysics have increased substantially. This has led to an expansion in the potential scope of evolutionary analyzes and a growing awareness of the limitations of standard empirical models. Proteins are under selection for traits – function, foldability, stability, solubility – that depend on a complex network of interacting amino acids. These forms of selection induce epistatic interactions or coevolution) among sites in the protein, resulting in substantial effects on the evolutionary process (*7–12*). Models that ignore this epistasis can seriously compromise evolutionary analyzes by misrepresenting the frequency and time dependence of convergence and homoplasy (*13*). Empirical models can be modified to allow the substitution process to vary among sites (*14, 15*) and over time (*6, 16–20*), but information available from sequences to obtain accurate site- and time-dependent substitution rates is fundamentally limited. Efforts to use protein structure to predict substitution rates (*21, 22*) are compromised by our lack of understanding of the relationship between protein sequence, protein properties, and organismal fitness, and our inability to predict the effect of mutations as differences accumulate.

The development of more accurate and powerful models of protein evolution depends on our ability to represent the process of molecular evolution at a mechanistic level, ideally enabling us to calculate substitution rates based on the protein’s sequence and biophysical properties. Our purpose here is to develop from first principles a theory of how proteins evolve and how substitution rates are determined. We approach the problem by building a conceptual framework to translate protein evolution into the formalisms of statistical mechanics, demonstrating the primacy of sequence entropy. Using evolutionary simulations of model proteins, with fitness determined by thermodynamic stability, we demonstrate that substitution rates depend on how amino acid energy contributions fluctuate as the rest of the protein sequence evolves. We show that substitution rates can be predicted based on 1) the stability distributions at a site in the absence of selection on that site; and 2) the relative numbers of sequences with different protein stabilities; no other adjustable parameters, such as expected population size, are needed. This forms a mechanistic framework for constructing improved models of amino acid substitution rates.

## Results

### Site-specific stabilities and relative substitution rates

To develop a mechanical theory of the evolutionary process, we consider the relationship between protein stability and substitution rates at a site. The stability Ξ(**X**) of a protein sequence **X** = {*x*_1_, *x*_2_, *x*_3_… *x_n_*} was defined as the negative of the free energy difference between the sequence in the native structure and in the ensemble of possible alternative structures, so that more positive values indicate greater stability. The Malthusian fitness *m*(**X**) was set equal to the fraction of such sequences that would be folded in a pre-specified native conformation at thermodynamic equilibrium (Equation (2, Methods) (*11, 23, 24*). Thus, increases in stability lead to increases in fitness.

To understand how the rest of the protein influences the substitution rate at individual sites, we focus our attention on an amino acid α at a specific focal site *k*, and partition the stability into Ξ(**X**) = ξ_*k*,α_(**X**_∌*k*_) + ξ_*k*,Bath_(**X**_∌*k*_). The first term, ξ_*k*,α_(**X**_∌*k*_), is the site-specific stability contribution due to interactions (in both the folded and unfolded states) between *α* at site *k* and the amino acids at all other sites excluding *k*. The second term, ξ_*k*,Bath_(**X**_∌*k*_), is the ‘background’ contribution resulting from interactions among amino acids at sites excluding the focal site. Because only a small fraction of contacts involve site *k*, we assume that the site-specific stability contribution is small relative to the background contribution, so that this second term fulfills the role of the ‘thermal bath’ in statistical physics.

This statistical mechanics formalism can now be applied to modeling the amino acid substitution rate. Consider *Q*_*k*,α→β_(**X**_∌*k*_), the instantaneous rate of an α to β substitution at site *k*, equal to the mutation rate times the fixation probability. The fixation probability depends on the difference in fitnesses Δ*m_*k*α→β_*, which is a function of the initial stability Ξ(**X**) and the stability of the mutant Ξ(**X**′) = Ξ(**X**) + ΔΞ_*k*,α→β_(**X**_∌*k*_). We can simplify the situation by noting that sequences from real proteins, as well as proteins from evolutionary simulations under selection for thermostability, tend to have a narrow range of stability values(*23, 25–28*). This stability range occurs where the decreasing effectiveness of selection for greater stability is balanced by destabilizing mutations fixed by genetic drift. The precise value depends on a variety of factors such as temperature, effective population size, sequence length and protein function. As long as these factors are approximately constant, we can assume that a given protein will evolve to the mean of this narrow range 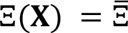. If 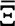 is a known constant, calculating the fixation probability requires only the difference in site-specific stabilities ΔΞ_*k*,α→β_(**X**_∌*k*_) = ξ_*k*,β_(**X**_∌*k*_) − ξ_*k*,α_(**X**_∌*k*_); the bath component, ξ_*k*,Bath_(**X**_∌*k*_) is independent of the amino acid at focal site *k*, and is therefore unchanged by the substitution.

From this perspective, the key distribution determining the substitution rate from α to β is ρ_*k*,α_(ξ_*k*,α_, ξ_*k*,β_), the joint probability density of ξ_*k*,α_(**X**_∌*k*_) and ξ_*k*,β_(**X**_∌*k*_) given that amino acid α is resident at site *k*, integrating over the distributions of amino acids at other locations. The distribution depends on which amino acid occupies position *k* because that amino acid will have affected the evolution in the rest of the protein; in this case, ξ_*k*,β_ is the local stability contribution that would result if β were to replace α at that site with no other changes in the sequence. For simplicity, we will consider that the rest of the protein sequence has evolved sufficiently that ρ_*k*,α_(ξ_*k*,α_, ξ_*k*,β_) has reached a stationary distribution; the effect of a breakdown in this assumption will be considered below.

To help visualize these distributions, and evaluate our theoretical results, we modeled the evolution of real proteins using the simulated evolution of a 300-residue protein under selection for thermodynamic stability. This model is not meant to make quantitative predictions in particular cases. Instead, it is meant to predict general characteristics of evolutionary behavior for proteins that require the native confirmation to carry out some critical biological function, and has demonstrated its ability to reproduce fundamental aspects of the evolutionary process (*11, 23, 24*). By using a simple pair-contact model of protein thermodynamics, we were able to perform replicate simulations over long periods of evolutionary time, corresponding to approximately 5 billion years given typical substitution rates.

We grouped sites with similar substitution patterns into four different site classes, where class 1 is the most exposed and 4 is the most buried. Figures 1A-D shows the observed joint probability distributions of these site classes for glutamic acid and lysine, as well as the stability distributions when substitutions between these two amino acids occurred. Figures 1E-H shows these distributions for four different pairs of amino acids in site class 3. There are wide ranges of values for ξ_*k*,α_ and ξ_*k*,β_, consistent with earlier results demonstrating fluctuating selective pressures at sites due to substitutions elsewhere in the protein (*11*). The distributions of ξ_*k*,α_ and ξ_*k*,β_ strongly depend on the resident amino acid. In particular, the potential contribution of an amino acid to the protein stability tends to be greater when that amino acid is resident at a site, a phenomenon we previously named the ‘evolutionary Stokes shift’ (*11*). The amount of this increase appears to be correlated with the observed variance in ξ_*k*,α_.

**Figure 1:**
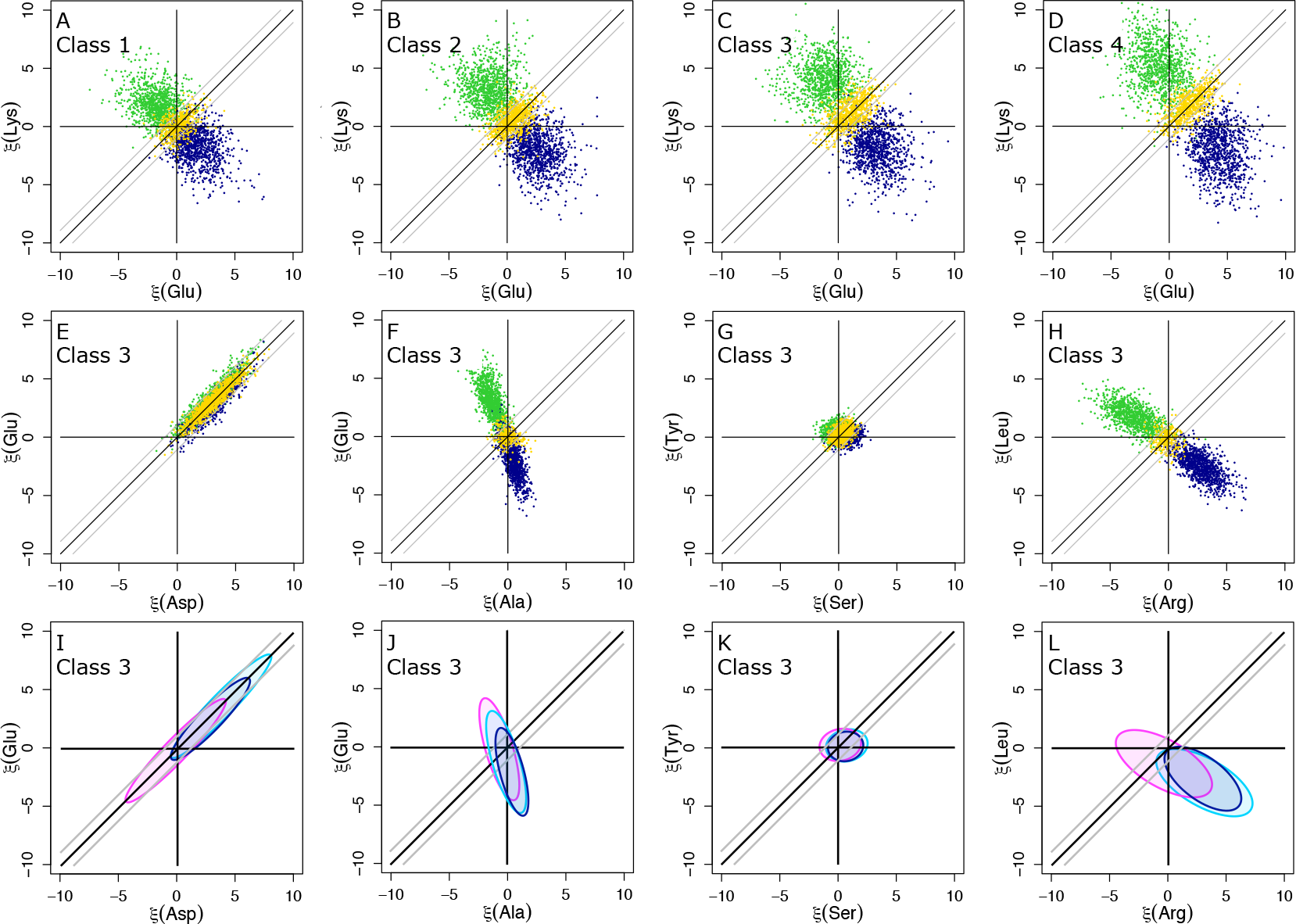
A-H) Relative local contributions to stability for pair of amino acids in different site classes (A-D) or different pairs of amino acids in the same site class (E-H). Points were sampled either when the amino acid in the abscissa is resident (blue), when the amino acid in the ordinate is resident (green), or during transitions between the two (yellow). I-L: Distributions of local contributions to stability in reference state when the non-interacting null amino acid was present ρ_*k*,∅_(ξ_*k*,α_,ξ_*k*,β_), megenta), when the amino acid in the abscissa was present as predicted using Equation (1 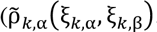, cyan), or as observed *ρ*_*k*,α_(ξ_*k*,α_, ξ_*k*,β_), blue).

Rapidly evolving sites with few selective constraints tend to have compact distributions with smaller variances in ξ_*k*,α_ and ξ_*k*,β_ than slowly evolving sites (Figures 1A-D). Distributions for physicochemically similar amino acids (e.g., aspartic acid versus glutamic acid, Figure 1E) appear highly correlated, while those for dissimilar amino acids (e.g., arginine versus leucine, Figure 1H) seem anti-correlated. This is because background sequences that confer a high site-specific stability on aspartic acid tend to do the same for the highly similar glutamic acid, while background sequences that stabilize arginine tend to destabilize the dissimilar leucine (*29*). A non-resident amino acid is generally stabilized if the distributions are correlated (e.g. glutamic acid when aspartic acid is present, Figure 1E), but destabilized if the distributions are anti-correlated (e.g. glutamic acid when alanine is present, Figure 1F).

To determine whether substitution rates can be predicted from ρ_*k*,α_(ξ_*k*,α_, ξ_*k*,β_), class-specific stability distributions were modeled with the best fitting bivariate normal distribution for each pair of amino acids 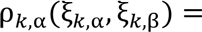 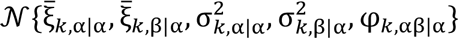. The expected substitution rates between each pair were then estimated by numerical integration over these distributions using Kimura’s formula for the probability of fixation (*30–32*) (see Equation (3, Methods). There is extremely good agreement between expected substitution rates derived from this approximation and substitution rates obtained by counting substitutions that occurred during the simulations (Figure 2). This validates the utility of the bivariate normal approximation and the assumption that variation in Ξ(**X**) has little effect on substitution rates.

**Figure 2:**
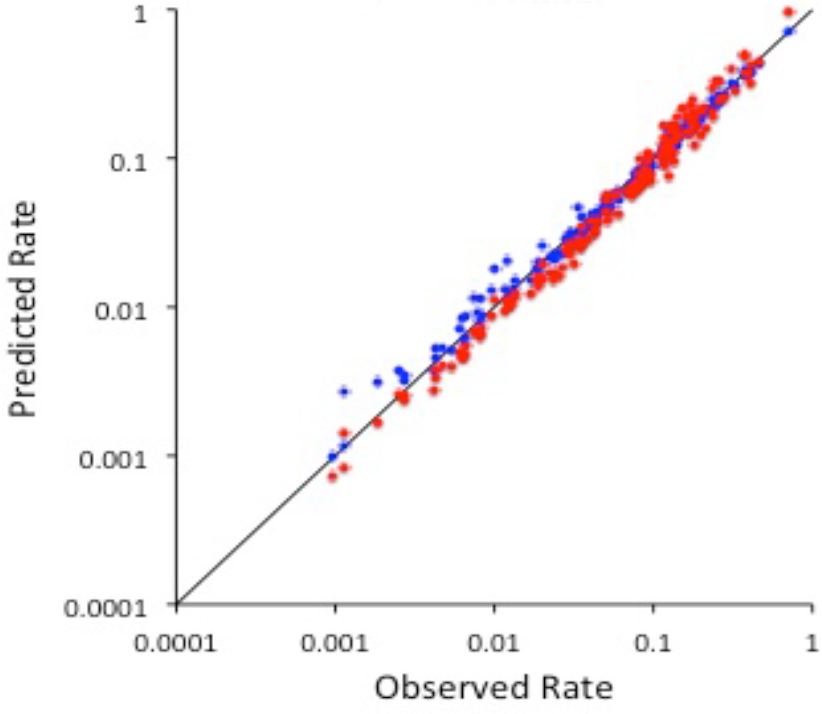
Comparison of observed and predicted substitution rates. Blue: predicted substitution rates calculated by integrating over *ρ*_*k,α*_(*ξ_k,α_*, *ξ_k,β_*), Red: Predicted substitution rates calculated using transition state theory (Equation (7), which assumes only near-neutral substitutions occur.

A striking feature of Figure 1 is the strong tendency for substitutions to occur in the overlap region between ρ_*k*,α_(ξ_*k*,α_, ξ_*k*,β_) and ρ_*k*,β_(ξ_*k*,α_, ξ_*k*,β_), centred on the diagonal ΔΞ_*k*,α→β_ = ξ_*k*,β_ − ξ_*k*,α_ = 0 where substitutions are neutral. This suggests the possible applicability of transition state theory (TST), a method for predicting the rate of chemical reactions (*33*). In TST, the reaction rate is given by the fraction of reactants in a ‘transition state’ in which the energies of reactant and product are approximately equal, times the rate of conversion from transition state to products. Adapting this theory, we model the substitution rate as equal to the fraction of joint stabilities for which the fitness of wild type and mutant are approximately equal, times the rate of substitution under neutral conditions.

The probability that the background sequence results in nearly equal fitnesses between α and β at site k was estimated as the density ρ_*k*,α_(ξ_*k*,α_, ξ_*k*,β_) integrated along the neutral line ξ_*k*,α_ = ξ_*k*,β_, multiplied by the width of the neutral zone on both sides of the neutral line, 2ε, the region in which the effect of selection is small. The neutral substitution rate is equal to the mutation rate *v*_α→β_, allowing us to write a closed-form expression for the average substitution rate (Equation (7, Methods).

As described in the Methods section, the extent of the neutral zone, ε, can be naturally defined by the falloff in the number of sequences with greater stabilities. Because the stability values Ξ for folded proteins represent the far tail of a distribution dominated by unstable sequences, we modeled Ω(Ξ), the number of sequences with stability Ξ, as an exponential Ω(Ξ) ∝ exp(−γΞ), where γ characterizes the decrease in number of sequence with increasing stability; thus, the bias of the drift effect does not depend on 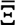. The scale of the neutral zone is given by 1/γ, which is equal to the range of stabilities at which the fitness changes by less than 1/4*N_e_* regardless of population size (see Methods). To calculate substitution rates, we estimated γ = 1.26 (kcal mol-1)-1 based on the relative numbers of destabilizing and stabilizing mutations, yielding ε = 0.79 kcal mol-1. Notably, because this calculation considers only neutral substitutions, it produces strikingly accurate predictions (Figure 2) without the need for Kimura’s formula.

### The equilibrium distributions of site-specific stabilities: The mechanism behind the evolutionary ‘Stokes Shift’

It appears that the rate of amino acid substitutions is substantially determined by ρ_*k*,α_(ξ_*k*,α_, ξ_*k*,β_) in the regions where 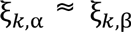. Our goal for the rest of the paper is to show the degree to which these distributions, and therefore substitution rates, can be explained using the principles of statistical mechanics.

As above, we assume that proteins evolve to a specific stability value 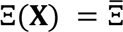. All sequences with stability 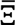 have, in our model, identical fitnesses, so none are preferred over another by selection. If evolution has had sufficient time to sample from the stationary distribution, the fraction of sequences with any property is proportional to 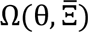, the number of sequences with property θ and stability 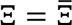. The log of this quantity, 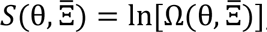, is the ‘sequence entropy’ of such sequences, analogous to thermodynamic entropy. Under these conditions, the probability of property θ is given by 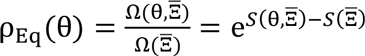, where 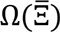 and 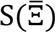 are the number of sequences with stability 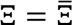 and the log of this quantity.

To calculate 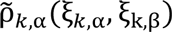, an estimate of ρ_*k*,α_(ξ_*k*,α_, ξ_*k*,β_), we first considered 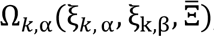, the number of sequences with stability 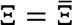, α resident at site *k*, and site-specific stability contributions ξ_*k*,α_ and ξ_*k*,β_ at that location. We approximated this number as the product of 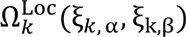, the number of amino acid arrangements resulting in the site-specific ξ_*k*,α_ and ξ_*k*,β_, times 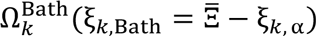. The latter term is the number of sequences furnishing the background stability required to complement the site-specific contribution furnished by ξ_*k*,α_, for a total stability equal to 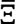. This calculation assumes independence of the bath and local contributions to total stability; although not strictly accurate (the relevant sites in the protein overlap), it is likely to be approximately true because the interactions involved are different.

We note that 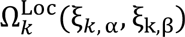 does not depend on selection, so to characterize it requires removing selection at site *k*. To do this we performed simulations with the focal site permanently occupied by a non-interacting amino acid, ∅, and with all other sites evolving freely. The resulting distributions ρ_*k*,∅_(ξ_*k*,α_,ξ_k,β_) are proportional to 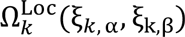, and represent the null distributions of local stability contributions that would occur if interactions between amino acids at site *k* and the rest of the protein did not affect the evolutionary dynamics. Because the number of possible sequences is immense, and because ξ_*k*,α_ and ξ_k,β_ are the result of many interactions, the central limit theorem suggests that ρ_*k*,∅_(ξ_*k*,α_,ξ_k,β_) can be approximated by a bivariate normal distribution 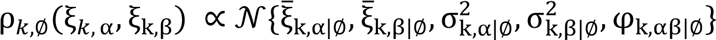. Interactions involving the focal amino acid represent a small fraction of total interactions, allowing us to approximate 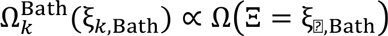. The normalized product of ρ_*k*,∅_(ξ_*k*,α_,ξ_k,β_) and the exponential 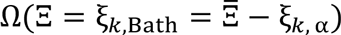 results in a shifted bivariate normal distribution 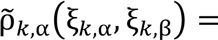 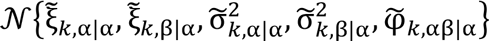 with

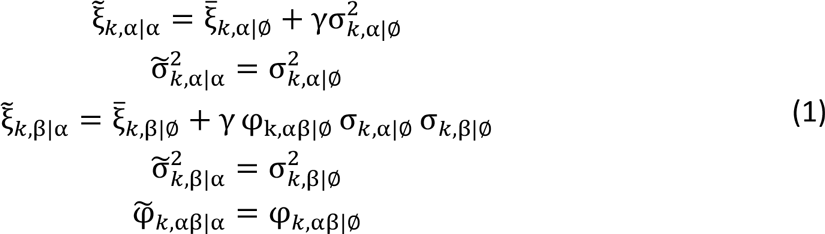

Selection in the presence of amino acid α at site *k* shifts the average local contribution to stability by an amount 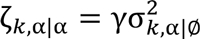 compared to its contribution to stability in the absence of interactions; this stabilization can be viewed as the basis for the evolutionary Stokes shift. The mechanism for the shift is the large increase in sequence entropy gained from a decrease in ξ_*k*,Bath_, combined with the trade-off between ξ_*k*,α_ and 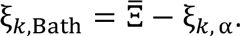.

The fit between estimated equilibrium values of 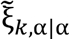 for each amino acid and values of 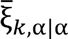 calculated directly from the simulations is surprisingly good given the approximations made (Figure 3A). The entropic stabilization as a function of 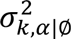 is linear (correlation coefficient 0.857; Figures 3B) as predicted by Equation (1. The slope of 1.00 (kcal mol^−1^)^−1^ (95% CI: 0.87 − 1.14) is close to the expected value of γ = 1.26 (kcal mol^−1^)^−1^, confirming the trends evident in Figure 1. The observed entropic stabilization is smaller than predicted for the two largest shifts in the slowest rate class, involving the negatively charged aspartic acid and glutamic acid. Earlier work demonstrated that equilibration for the most buried states can be extremely slow(*11*), and these outliers may represent cases where the protein has had insufficient time to adjust to the presence of the new amino acid.

The key result here is that the magnitude of the entropic stabilization that drives the evolutionary Stokes shift depends only on the number of protein sequences with given protein stabilities and on the underlying distributions of interactions in the absence of selection: the effect can be understood purely in terms of biophysics and sequence entropy.

**Figure 3:**
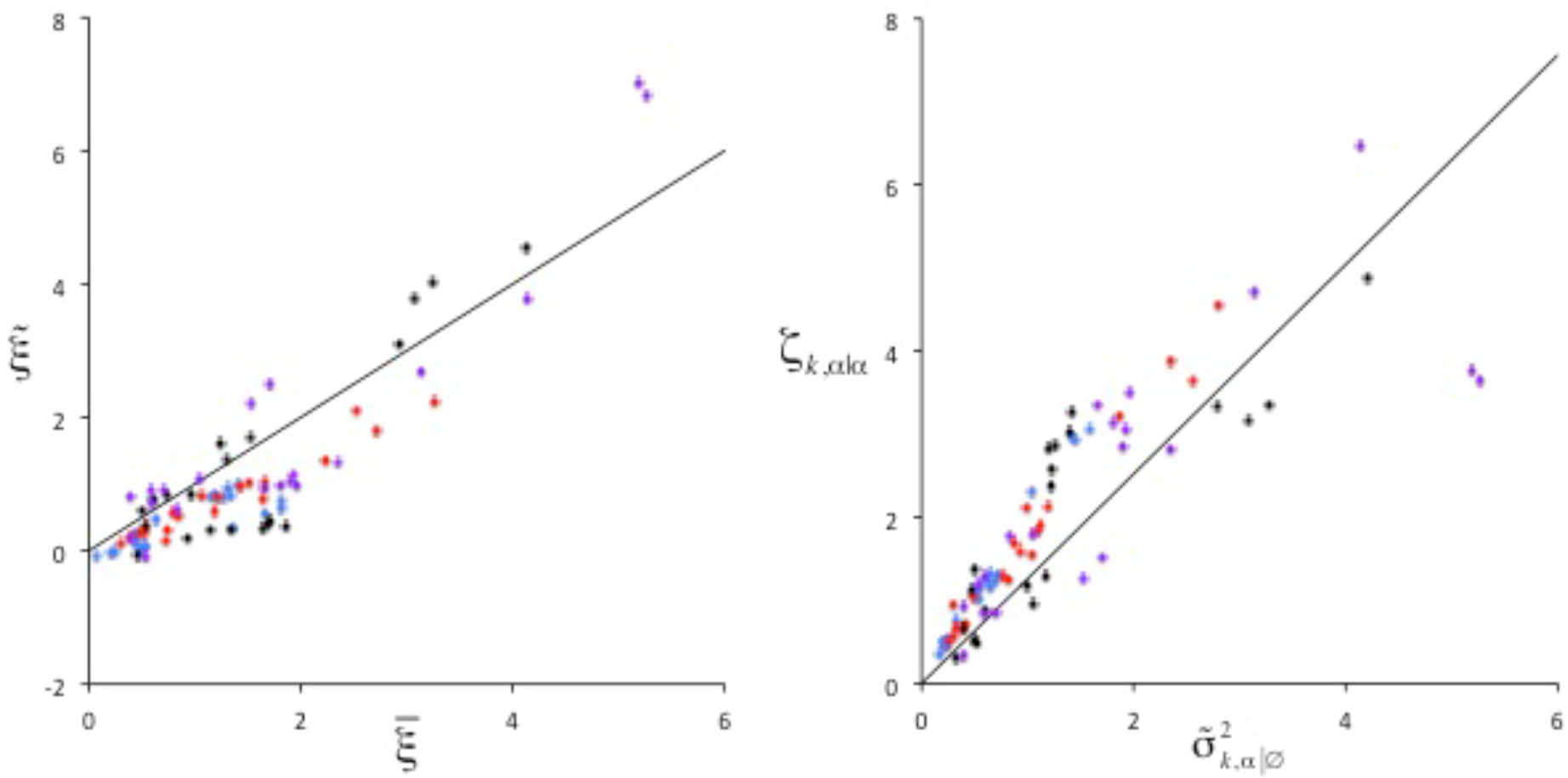
Accuracy of site-specific stability and evolutionary Stokes shift predictions. A) Estimated values of 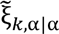 versus observed values 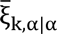 for all four site rate classes (from most exposed to most buried: Class 1, blue; Class 2, red; Class 3, black; and Class 4, purple). B) The linear relationship between the observed evolutionary Stokes shift and the variance in amino acid-specific stability contributions in the absence of selection on the site. The lines shown are theoretical predictions with gamma = 1.26.

The predicted and observed distributions of ρ_*k*,α_(ξ_*k*,α_, ξ_*k*,β_) are shown in Figures 1I-L. The values of ξ_*k*,β_ are shifted by an amount 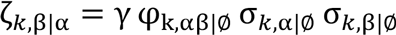 to either higher or lower values depending on the physicochemical similarities between the amino acids. From Equation (1 we can see that the realized evolutionary ‘Stokes shift’ after a substitution, the expected average difference in stability before and after the protein adjusts to the new resident amino acid, is equal to 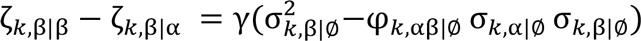. The full entropic stabilization is reduced by ζ_*k*,β|α_, which can be viewed as the average amount of preadaptation (or lack thereof) to amino acid β caused by the residency of amino acid α. As w with the average entropic stabilization, the realized evolutionary Stokes shifts *depend deterministically on the site-specific stability distributions in the absence of selection with no adjustable parameters*. Substitution rates estimated with the TST approximation (Equation (7) using the site-specific stabilities calculated from Equation (1 are remarkably accurate for all four site classes and over four orders of magnitude of rate variation (Figure 4).

**Figure 4:**
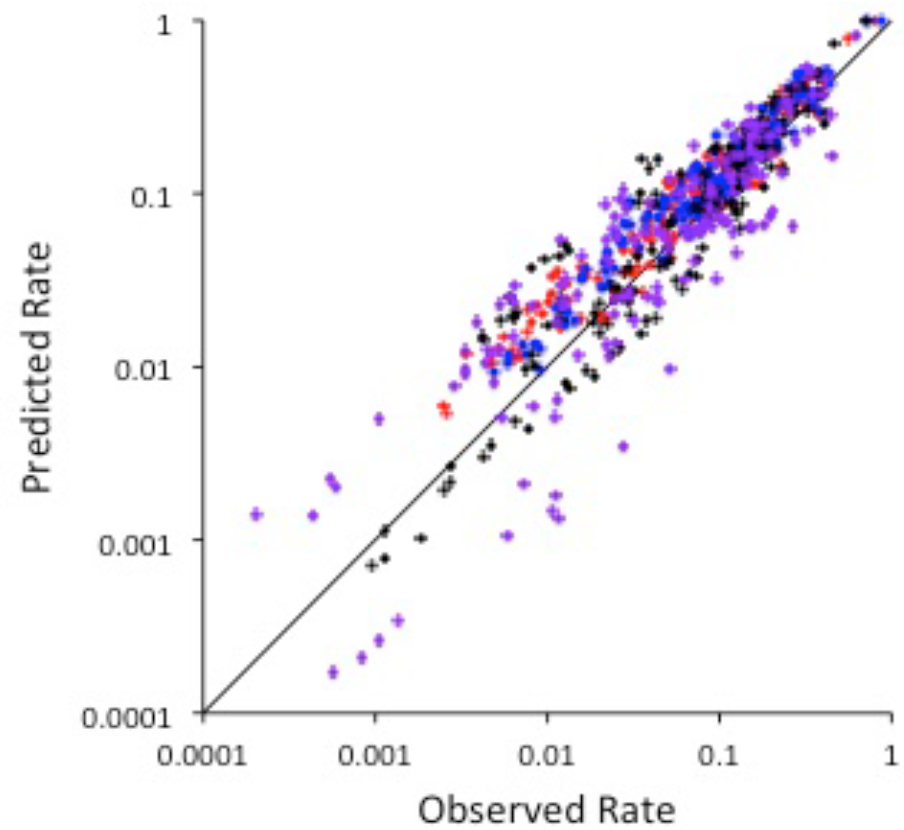
Predicted and observed values of substitution rates based on transition state theory. Rates were computed using estimated values compared with observed values for all four classes (Class 1, blue; Class 2, red; Class 3, black; Class 4, purple).

## Discussion

The understanding of evolutionary mechanics developed here represents a fundamental shift inhow we conceptualize the process of amino acid substitution. Although stability is approximately constant, the way this stability is partitioned among various interactions fluctuates as a protein evolves. In particular, the contribution that a resident amino acid at a site makes to the stability of the protein, as well as the contribution a non-resident amino acid would make if substituted in, will fluctuate. Occasionally, these fluctuations lead to approximately equal stabilities for a pair of amino acids so that substitutions from one to the other are nearly neutral. The frequencies of these nearly neutral states then determine the relative substitution rates. The fluctuations in stabilizing contributions of different amino acids at a site are not superfluous or unwanted complications in the construction of substitution models, but rather are central to the substitution process. In evolutionary theory it is common to evoke the idea of a fixed ‘adaptive landscape’, but for a single amino acid position a more appropriate analogy may be a fluctuating adaptive seascape; the site explores the space of possible amino acids by moving along fluctuating local contours in the context of approximately constant overall fitness.

By developing a statistical mechanics view of protein evolution, the evolutionary Stokes shift can be seen as a direct consequence of sequence entropy. Increases in the stabilizing contributions of an amino acid occupying a given site reduce the amount of stabilization required by the rest of the sequence, increasing the number of sequences that can contribute this reduced stability. Our theoretical analysis of the balance between the number of states available to the system (the amino acid at the focal site and its interactions) and the ‘bath’ (the rest of the sequence) yields an expectation that the relative magnitude of entropic stabilization of an amino acid at a site is proportional to the variance of the underlying null site-specific stability distribution. Furthermore, the stabilization of all amino acids at all sites are scaled by a protein-wide proportionality constant determined by the decline in the number of available sequences as protein stability increases. Thus, surprisingly, the strength of selection and the effective population size do not affect the evolutionary Stokes shift or substitution rates if the protein is in a steady state(*34, 35*). Thus, although our evolutionary mechanics theory fully incorporates population genetics theory and Kimura’s equation for the probability of a substitution, if the system is near equilibrium we do not need Kimura’s formula to predict and explain substitution rates among amino acids.

Correlations in the fluctuations between amino acids with similar physicochemical properties increase the probability of near-neutrality, providing a mechanistic explanation for higher rates of conservative change, a general phenomenon rationalized by Fisher with his geometric argument(*36*). The probability of occupying the neutral zone is lower at interior sites, where the multiplicity of interactions with the focal site increase the distance between ξ_*k*,α_ and ξ_*k*,β_ and correspondingly higher at surface sites; this is consistent with observed slower internal (buried) than external (surface) substitution rates.

For dissimilar amino acids, the probability of achieving the near-neutrality required for a substitution can be unlikely. However, if such a substitution occurs the protein will subsequently evolve to sequences that partition a larger stability contribution to the newly resident amino acid, causing an increased affinity for this residue. This increased affinity is what we have called the evolutionary Stokes shift. This evolutionary mechanism can be fully reversible, as is in our evolutionary simulations, with the reversibility coming from the similarity in the processes of moving into and away from the neutral zone (*11*). These processes, called ‘contingency’ and ‘entrenchment’ by Plotkin and colleagues (*12*), are mirrors of each other, so that if the substitution were reversed the dissipation process, played backwards, would have the same statistical properties as the pre-adaptation process played forwards. Where previously we might have assumed that the amino acid found at the site had adapted to the requirements of the site, the site may have instead adapted to the resident amino acid.

The fluctuations and the relaxation of the protein are explicitly time-dependent. Here we addressed only the theoretical equilibrium predictions and the result of simulations that were designed to be near equilibrium. This neglect of this time dependence may explain some of the errors in the predicted Stokes shift for charged residues in buried sites. Individual sites at specific time points might be further constrained by conserved neighboring sites in the structure as well as the conserved structural context of their interactions with those sites. Such effects may influence the time-dependent probability of back mutations as well as subsequent substitutions, an important topic for further investigation.

The simulations presented here also consider only the fitness effects of stability, but fitness is also usually determined by other effects such as interactions with substrates, ligands or other proteins. Such alternative fitness components will add additional constraints to the system, and may force non-neutral substitutions if outside selective pressures change. Previous analyzes indicated that when a substitution is compelled by an outside force, an evolutionary Stokes shift occurs in largely the same fashion, except that the process is no longer reversible (*11*). In this context, evolution can be seen as occurring in a ‘memory foam’ made up by the bath of interactions that occur among all sites other than the selected focal site.

In conclusion, the work described here sets up a theory of evolutionary mechanics, and demonstrates that this theory can be used to predict substitution rates from the basic properties of how amino acids interact. Although the current work is focused on fitness defined by the protein stability, we expect that other kinds of selection will fit well into this framework, either by defining a large nearly neutral landscape in their own right, or by constraining the stability-based nearly neutral network.

## Methods

### Simulations of protein evolution

The methods used to simulate protein evolution have been described previously (*11, 23, 24*). The free energy *G*(**X, r**) of a protein sequence **X** = *{*x*_1_, *x*_2_, *x*_3_*… *x_n_*} in conformation **r** was calculated by summing the pair-wise energies of amino acids in contact in that conformation, using the contact potentials derived by Miyazawa and Jernigan (*37*). We computed the free energy of folding Δ*G*_Folding_(**X**) by first determining the free energy of the sequence in a prechosen native state, the conformation of the 300-residue purple acid phosphatase, PDB 1QHW (*38*)). The energies of the unfolded states were assumed to follow a Gaussian distribution with parameters estimated by calculating the free energy of the sequence in an ensemble of 55 different structurally diverse protein structures. The energy of the unfolded state was then calculated by assuming a large set (10^160^) of possible unfolded structures with free energies drawn from that distribution. The free energy of folding Δ*G*_Folding_(**X**) was calculated as the difference between the two, and stability was Ξ(**X**) = −Δ*G*_Folding_(**X**). The Malthusian fitness of a sequence *m*(**X**) was defined as the fraction of that sequence that would be folded to the native state at equilibrium

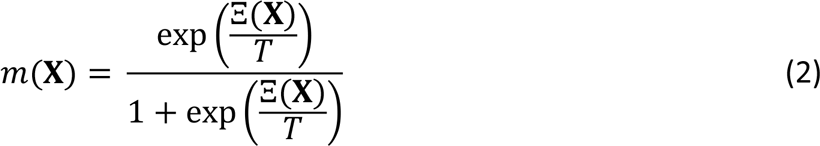

where *T* is the temperature in units of energy, 0.6 kcal mol^−1^.

Starting from a randomly chosen nucleotide sequence encoding a 300 amino-acid protein, we simulated evolution by considering in each step all possible nucleotide mutations with rates given by the K80 nucleotide model (κ = 2)(*39*). The fixation probability of each mutation was calculated based on the Kimura formula for diploid organisms (*30–32*),

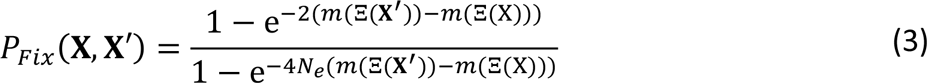

where **X** and **X′** are the sequences before and after the mutation, with the effective population size *N_e_* set to 10^6^. One substitution was chosen to be fixed at random with relative probabilities determined by the product of the mutation rates times the acceptance probabilities.

Sequence evolution was simulated for a sufficient number of generations such that the stability of the protein was roughly constant, representing mutation-drift selection balance. 100 such equilibrated proteins were chosen, and three longer simulations were performed using each these equilibrated proteins as initial starting sequences, for a total of 300 simulations. We simulated the evolution of each lineage for an evolutionary distance of approximately seven amino acid replacements per amino acid position.

### Grouping of sites

For ease of analysis, we divided the sites in the protein into four classes with similar substitution rates. Substitution matrices were calculated individually for each site; due to the length of the simulations, we had on average over 2000 substitutions at each site. We then clustered the sites based on the off-diagonal elements of the substitution matrices using K-means clustering (*40, 41*). The resulting clusters were approximately of equal size, and class membership strongly dependent on how buried or exposed the sites were in the native state (as indicated by number of contacts). We ranked the clusters by surface exposure, where class 1 is the most exposed and 4 is the most buried.

### Calculating the site-specific contribution to protein stability

The site-specific contribution ξ_*k*,α_(**X**_∌*k*_) of amino acid α at focal site *k* as a function of the amino acids **X**_∌*k*_ at all sites excluding *k* is equal to Ξ{*x*_1_, *x*_2_,*x*_3_… *x*_*k*−1_, α, *x*_*k*+1_… *x_n_*}, the stability when the focal site is occupied by α, minus Ξ{*x*_1_, *x*_2_,*x*_3_… *x*_*k*−1_, ∅, *x*_*k*+1_… *x_n_*}, the stability of a reference state when by a is replaced by a non-interacting amino acid ∅, while the rest of the sequence and thus all other interactions, are unchanged. The part of the stability unaffected by this replacement is represented by the ‘bath’ interactions ξ_*k*Bath_ (**X**_∌*k*_) so that Ξ(**X**) = ξ_*k*,α_(**X**_∌*k*_) + ξ_*k*Bath_ (**X**_∌*k*_).

### Calculating the substitution rate integrating over distributions of local contributions

The average rate for the substitution α → β at site *k*, *Q*_*k*,α→β_, is equal to the neutral substitution rate *v*_α→β_ times the average probability of fixation, which is a function of the stability of the protein before and after the substitution. The standard deviation of observed values of Ξ, 0.71 kcal mol^−1^, was small compared with the range of values of ξ_*k*,β_ (as shown in Figure 1), allowing us to represent the distribution Ξ by its average,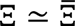 = 9.27 kcal mol^−1^. We assumed that the stability before the substitution was equal to 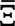 and after the substitution was 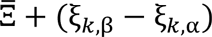. The average substitution rate was then estimated as

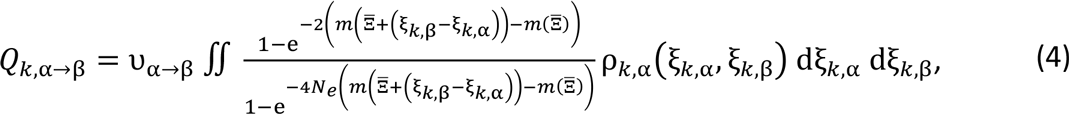

where ρ_*k*,α_(ξ_*k*,α_, ξ_*k*,β_) is the joint distribution of ξ_*k*,α_, and ξ_*k*,β_ observed when α occupies site *k*.

Based on the observations in Figure 1, we modeled ρ_*k*,α_(ξ_*k*,α_, ξ_*k*,β_) as a bivariate normal distribution of the form 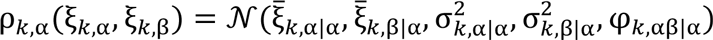, where the parameters are represented as explicitly depending on the amino acid occupying site *k*. These parameters were calculated directly from the evolutionary simulation, and Equation (4 was integrated numerically. The neutral substitution rate was calculated using the same K80 nucleotide model (κ = 2)(*39*) as used in the simulation, with all non-nonsense codons considered equally likely.

### Calculating the substitution rate integrating assuming only neutral substitutions

As observed in Figure 1, substitutions generally occur in a neutral region in which ΔΞ_*k*,α→β_ = ξ_*k*,α_ − ξ_*k*,β_ ≈ 0, so that

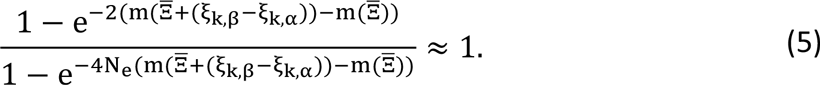

This condition is satisfied in a band of width 2ε centred on ξ_*k*,β_ = ξ_*k*,α_, where s represents the deviation from strict neutrality that is still sufficiently close for Equation (5 to be sufficiently accurate.

We can obtain a natural scale for ε by considering the concept of ‘free fitness’ Φ(Ξ) of the protein equal to Φ(Ξ) = *m*(Ξ) + *S*(Ξ)/4*N_e_* (*42, 43*). Free fitness, analogous to its thermodynamic equivalent ‘free energy’ where *T* is replaced by 4*N_e_*, encompasses the contributions of both fitness and sequence entropy in determining the distribution of states; evolutionary dynamics moves towards maximising this quantity. Assuming *S*(Ξ) = ln(Ω_0_ *e*^−γΞ^) where Ω_0_ is a constant, and noting that the system is at equilibrium with 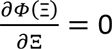 when 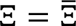, we can see that

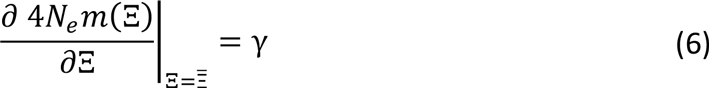

Thus, γ defines the rate of change of the population-weighted fitness 4*N_e_m*(Ξ) with stability. Alternatively, a change in stability of 1/γ corresponds to a unit change in the population-weighted fitness. In our calculations, we equated ε = 1/γ; the estimation of γ is described below. Note that this calculation demonstrates that ε is, surprisingly, independent of effective population size *N_e_*. This is a result of the balance between selection and mutational drift at equilibrium; for fixed effect of mutational drift, the degree of selection 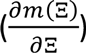 adjusts to changes in effective population size so that their product is constant(*34, 35*).

If we assume that ρ_*k*,α_(ξ_*k*,α_, ξ_*k*,β_) is broader than ε, and that Equation (5 is satisfied, Equation (4 becomes

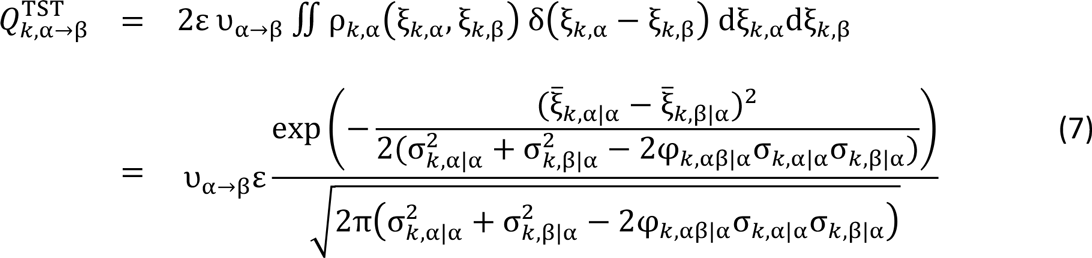

where δ(ξ_*k*,α_ − ξ_*k*,β_)is the Dirac delta function.

For highly similar amino acids the entire distribution of ρ_*k*,α_(ξ_*k*,α_, ξ_*k*,β_) may be contained in a region significantly narrower than the neutral zone, resulting in an overestimation of *Q*_*k*,α→β_ > *v*_α→β_. For this reason, the estimated rate was capped at the neutral rate *v*_α→β_.

### Characterising the bath state distribution

As described above, we assume that the number of protein sequences with a given value of Ξ in the range of interest around 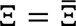 is approximately exponential Ω(Ξ)~ *e*^−γΞ^. To estimate γ, we consider the distribution of changes in stability resulting from random mutations, ρ_mut_(ΔΞ). The average change in stability 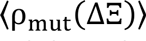 is negative due to the greater number of sequences coding for proteins with lower stability. This suggests that if we correct for the dependence of Ω on Ξ by multiplying ρ_mut_(ΔΞ) by *e*^γΔΞ^, this bias would disappear. We adjusted γ so that 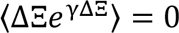 where the average was over all possible mutations during the simulations, yielding γ = 1.26 (kcal mol^−1^)^−1^.

